# Intranasal sertraline for the investigation of nose-to-brain delivery to mitigate systemic exposure

**DOI:** 10.1101/2025.08.10.669549

**Authors:** Shoshana C. Williams, Travis Lantz, Vanessa Doulames, Alakesh Alakesh, Daniel Ramos Mejia, Carolyn K. Jons, Noah Eckman, Eric A. Appel

## Abstract

Antenatal depression, or depression during pregnancy, is a common psychiatric disorder and poses significant risks to both the mother and the fetus. Despite these risks, it is frequently left untreated due to fears of side effects caused by antidepressant medications which cross through the placental barrier. It is therefore desirable to develop formulation strategies to mitigate systemic exposure to relevant drug molecules while maintaining their psychotropic efficacy. In this work, we develop formulations of sertraline, a common antidepressant, to target delivery to the brain through intranasal administration. Formulation engineering enables successful solubilization of sertraline at high concentrations and our lead formulation remains stable at room temperature for months. Using mice, we compare sertraline biodistribution following intranasal administration and standard oral administration. Intranasal administration of our drug product candidate provides comparable brain exposure at half the dose compared to oral treatment and lowers the maximum plasma exposure. These findings suggest that intranasal administration may provide selectivity for drug exposure in the central nervous system over systemic exposure.

## INTRODUCTION

Intranasal drug delivery has gained attention in recent decades, owing to its distinct advantages in several different indications. Initially, intranasal delivery was employed for localized drug delivery for indications such as allergic rhinitis or nasal polyposis,^1–4^ but it has since been implemented to treat systemic and neurological conditions.^5–8^ Intranasal administration offers large benefits for nose-to-brain delivery in particular, as it allows drugs to bypass the blood-brain-barrier (BBB) by shuttling therapeutics via the olfactory and trigeminal nerves.^9–11^ Intranasally administered drug products have been commercially implemented in the treatment of central nervous system (CNS) diseases such as depression, migraines, and opiate overdose to facilitate simple and rapid delivery of the pharmacological agent to the brain.^12–15^

While much research has extolled the benefits of nose-to-brain drug delivery,^6,9,16–20^ little work has focused directly on the question of selectivity for exposure in the brain over systemic exposure, or whether a drug can be targeted to the brain while diminishing systemic exposure. Intranasal administration avoids first-pass metabolism and has the potential to facilitate lower total dosages;^21^ however, the nasal epithelium is also highly vascularized, leading some amount of drug to be absorbed systemically.^22,23^ The question of selectivity is underexplored, although it is thought to depend on the specific formulation and delivery device.^8,10,24,25^

A selective nose-to-brain delivery approach would be invaluable in disease applications where systemic exposure would be undesirable due to potential side effects. For instance, antenatal depression, or depression during pregnancy, often goes untreated due to fears of potential side effects for the fetus. This disorder is estimated to affect 8.5% of American pregnant women,^26^ and is associated with severe adverse effects for both mother and fetus, including increased risk of preeclampsia, edema, hemorrhage, preterm birth and more (Figure 1A).^26^ Antenatal depression in the mother can also have a persistent adverse effect on the behavioral, neurological, and emotional development of the child (Figure 1A).^26^ Despite the serious risks it poses to mothers and infants, the disease is often left unchecked because to date, no psychotropic has been shown to be risk-free during pregnancy.^27^

**Figure 1.**
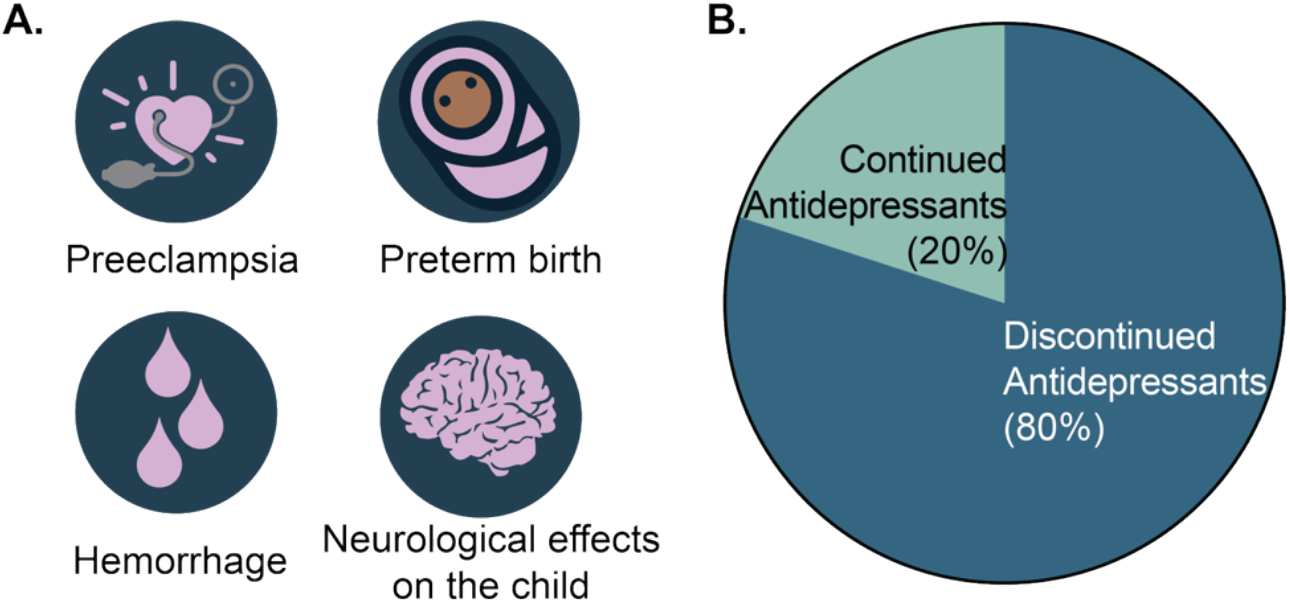
**A**. Complications of antenatal depression. **B**. Pie chart depicting the rate of antidepressant discontinuation among pregnant patients.

Common classes of antidepressant medications, including selective serotonin reuptake inhibitors (SSRIs), are considered among the safest antidepressants and are sometimes used during pregnancy, but each option is associated with potential complications for the fetus. Postnatal adaptation syndrome occurs in up to 30% of neonates exposed to antidepressants in late pregnancy.^28^ Due to this risk, as many as 80% of patients choose to discontinue antidepressant use upon becoming pregnant (Figure 1B).^29,30^ Strikingly, 68% of these patients experience relapse during their pregnancy,^31^ and they have a 52% increase in their risk of experiencing a psychiatric emergency during their pregnancy that results in a hospital visit compared to those who continued antidepressant use.^30^ It is therefore desirable to develop an antidepressant drug product capable of maintaining therapeutic efficacy while mitigating risks during pregnancy.

The most common antidepressant taken by non-pregnant reproductive-aged women is the generic drug sertraline, an SSRI used by 3.3% of this population.^32^ Sertraline is therefore an attractive candidate for reformulation to minimize risk during pregnancy. It is thought that the fetal risk stems from exposure to the drug in the amniotic sac,^33^ so it would be advantageous to explore formulations that could mitigate systemic, and thereby fetal, exposure to the drug. In this work we leverage polymeric excipients and formulation engineering to develop an intranasal sertraline drug product candidate and explore selectivity of sertraline exposure within the CNS in comparison to standard oral dosing.

## RESULTS AND DISCUSSION

### Development of a high-concentration formulation of sertraline for intranasal delivery

Sertraline is highly lipophilic, with a logP value of 5.51.^34^ This feature makes it an attractive candidate for intranasal delivery to the brain, as greater hydrophobicity enables preferential accumulation in the brain rather than bloodstream after oral administration.^35,36^ Although the hydrophobicity of sertraline may improve its uptake in the CNS,^37^ it also complicates the development of a liquid-phase aqueous formulations for nasal drops or a nasal spray as the solubility of sertraline in water has been reported to be only 3.8 mg/mL.^34^ To facilitate effective dosing, the concentration of sertraline in nasal drops should be at least 15 mg/mL, substantially higher than sertraline’s water solubility. To address this challenge, we evaluated various excipients for their ability to improve the solubility of the drug in an aqueous solution. All excipients tested are regarded as safe by the FDA and have been used in cosmetics and drug products.^38^ These excipients comprised various combinations of: β-cyclodextrin, γ-cyclodextrin, ethanol, isopropanol, propylene glycol, glycerol, dimethyl sulfoxide (DMSO), and 1-methyl-2-pyrrolidone (NMP). Previous work has demonstrated that encapsulation in β-cyclodextrin does not hinder the absorption or behavioral efficacy of orally-delivered sertraline, so this compound was not expected to interfere with an intranasal administration.^39^ Each formulation also incorporated a mucoadhesive polymer excipient, either pectin or hydroxypropyl methylcellulose (HPMC), to improve intranasal delivery. Such bioadhesives have previously been shown to improve the intranasal residence time of nasal drops.^11,40,41^ Aside from its activity as a bioadhesive, pectin also forms a gel in the presence of Ca^2+^ ions present in mucous, potentially enhancing its ability to retain a formulation in the nasal cavity over prolonged timeframes.^40,42^ Initial screening yielded two promising formulations, one incorporating each mucoadhesive polymer excipient (Table 1). These two formulations were further characterized.

**Table 1.** Composition of two formulations of sertraline.

|  | **Pectin Formulation** | **HPMC Formulation** |
| --- | --- | --- |
| Pectin (mg/mL) | 10 | 0 |
| HPMC (mg/mL) | 0 | 5 |
| β-Cyclodextrin/Sertraline complex (mg/mL) | 66.7 | 0 |
| Sertraline HCl | 0 | 20 |
| Glycerol (vol%) | 15 | 22.5 |
| NMP (vol%) | 6 | 9 |
| DMSO (vol%) | 2.5 | 3.8 |
| Water (vol%) | 76.5 | 64.8 |

Upon visual inspection, the pectin formulation appeared cloudy while the HPMC formulation was clear and colorless (Figure 2B-C). The cloudiness was believed to be due to the aggregation of pectin. The maximum sertraline concentration of each formulation was evaluated by monitoring absorbance at 260 nm. The pectin formulation had a maximum sertraline concentration of 22.6 mg/mL, while the HPMC formulation reached 35.1 mg/mL. Under the same conditions, the maximum drug concentration in water was found to be only 3.3 mg/mL (Figure 2D). Both formulations successfully enhanced the solubility of sertraline, with the HPMC formulation offering a tenfold increase over a standard aqueous formulation.

**Figure 2.**
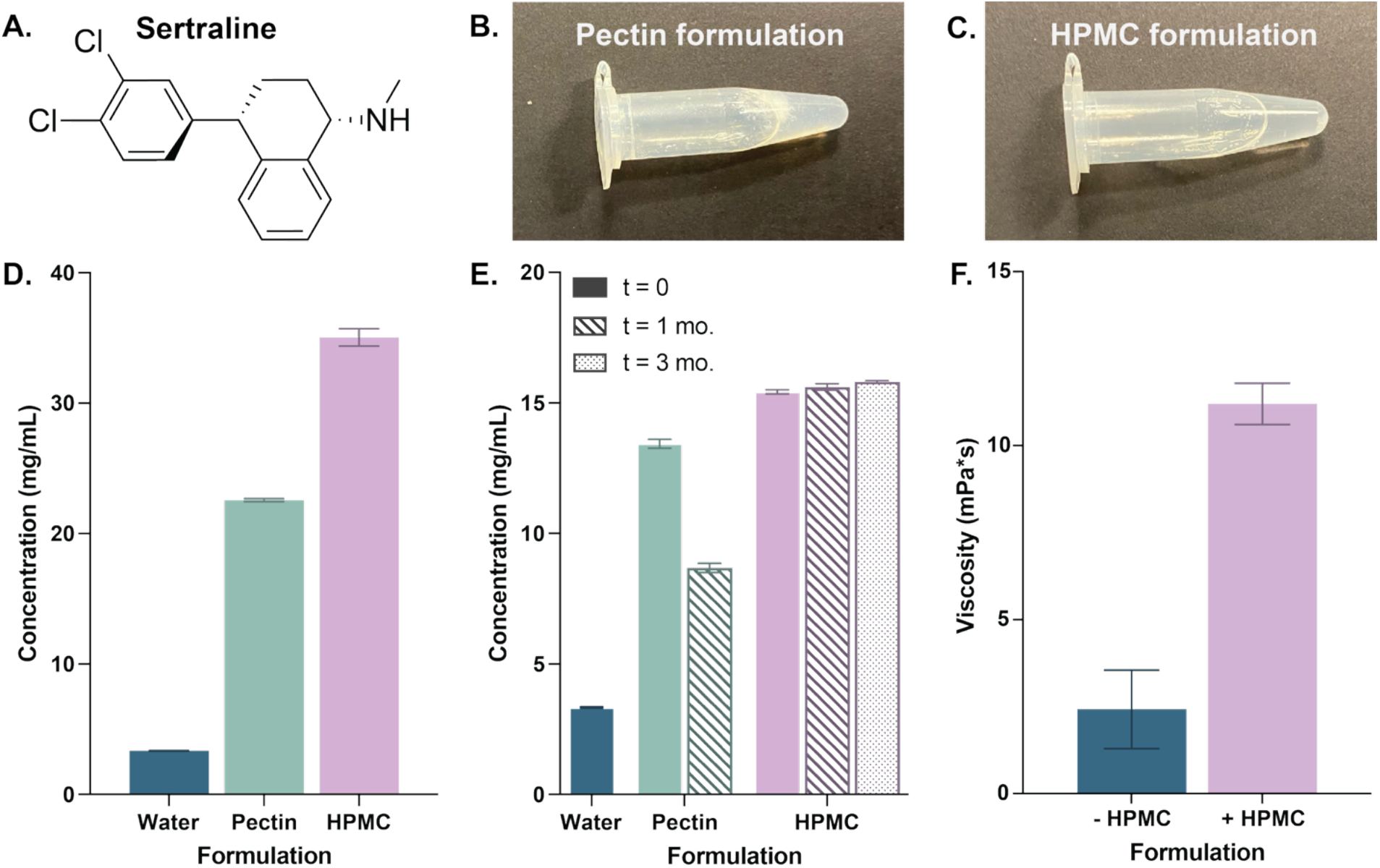
**A**. Chemical structure of sertraline. **B**. Photograph of sertraline formulation including pectin. **C**. Photograph of sertraline formulation including HPMC. **D**. Maximum concentration of sertraline in various formulations. **E**. Stability of sertraline in various formulations. **F**. Viscosity-modifying effect of HPMC. Error bars represent +/-SD.

These formulations were then evaluated for their stability over time. Both formulations were incubated at room temperature for several weeks, and their sertraline concentration was monitored over time to determine if the sertraline degraded or precipitated from solution. We found that the pectin formulation decreased in sertraline concentration after one month, while the HPMC formulation maintained stable sertraline concentrations for over three months (Figure 2E).

On account of its ability to solubilize a high concentration of sertraline in a stable fashion over months, the HPMC formulation was selected for further evaluation. The HPMC was chosen as a mucoadhesive and a viscosity modifier to reduce the likelihood of the formulation dripping down the throat or being inhaled into the lungs after intranasal administration. To characterize its effect, the viscosity of the formulation was measured in the presence and absence of HPMC (Figure 2F). It was found that the HPMC increased the viscosity of the solution from 2.4 to 11.2 mPa*s, indicating its efficacy in increasing viscosity without impacting its potential use as nasal drops or spray.^43,44^

### *In vivo* evaluation of intranasal sertraline

Due to its promising *in vitro* performance, the HPMC formulation was further evaluated *in vivo* in mice. Animals were dosed orally with 0.4 mg sertraline or intranasally with 0.2 mg sertraline. Animals were sacrificed at 15 min, 30 min, 2 h, or 4 h, with *n* = 9-11 animals per group per timepoint. The concentrations of sertraline in the plasma and in the brain were recorded for each animal (Figure 3A-B; Tables S1-4). For both plasma and brain tissue, the intranasally-treated animals showed higher sertraline concentrations at the initial timepoint of 15 min, as the drug likely had not yet been absorbed via the oral route. The concentration remained steadier over the course of the experiment for the intranasally-treated animals compared to the oral group, with the starkest differences between groups observed in the plasma concentrations.

**Figure 3.**
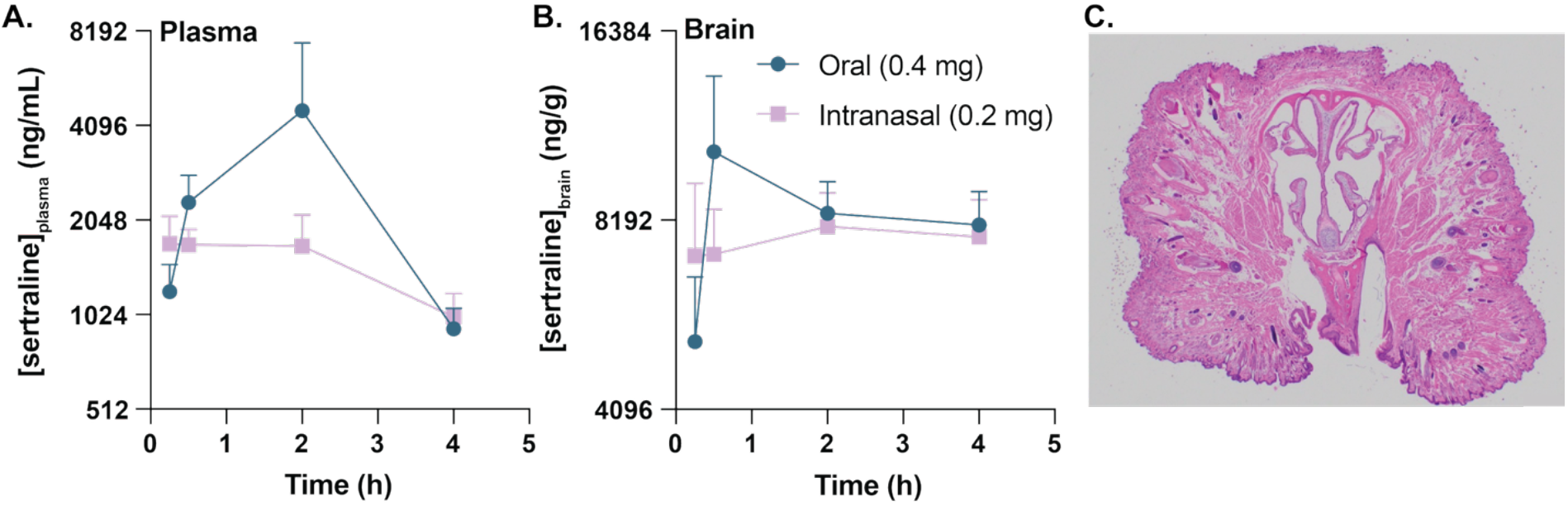
**A**. Plasma and **B**. brain concentration of sertraline after a single administration. Error bars represent SEM. **C**. Representative histological image of nasal passage of a mouse treated with intranasal sertraline, sacrificed 4h after administration.

To evaluate the safety of the intranasal administration of sertraline, histology was used to analyze tissue samples from the nasal epithelium, lung, and trachea for three animals sacrificed 4 h after treatment with intranasal sertraline. The results were evaluated for signs of acute toxicity and compared to corresponding samples from mice treated with oral sertraline. No discernible toxicity was observed in the samples from the intranasally-treated mice, suggesting that the formulation is well-tolerated in intranasal administration (Figure 3C; Table S5).

To analyze the results of the pharmacokinetic data, the area under the curve (AUC) of the brain concentration of sertraline was calculated for the population of mice (Figure 4A). No difference could be observed based on route of administration. This result is promising, given that the intranasally-treated animals received half the dose, compared to the orally treated mice. This result demonstrates that the lower intranasal dose is more efficient than oral, as it provides comparable brain exposure at a lower dosage.

**Figure 4.**
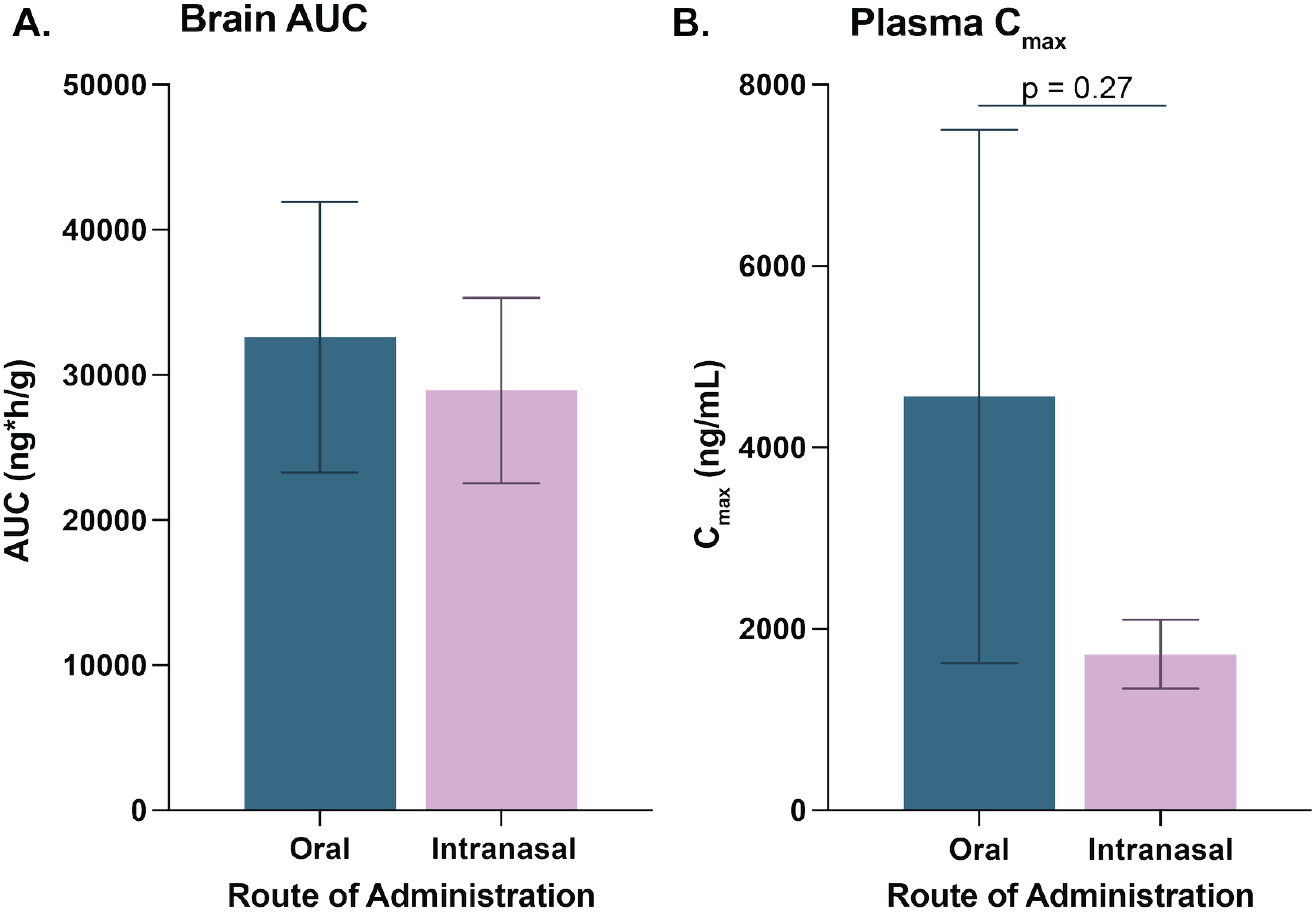
**A**. Area under the curve for brain concentration of sertraline +/-SEM. **B**. Maximum concentration of sertraline in plasma +/-SEM.

The maximum concentration (Cmax) present in the plasma was also calculated for the population of mice (Figure 4B). For the orally treated group, Cmax = 4700 ng/mL, while Cmax = 1700 ng/mL for the intranasally-treated group. This difference was not statistically significant (p = 0.27), owing to large variation among animals, particularly within the orally treated group; however, the results suggest that intranasal administration may diminish Cmax and reduce variability between animals.

## CONCLUSION AND OUTLOOK

This work investigated the potential of an intranasal administration of sertraline to provide selectivity and deliver the drug to the brain while mitigating systemic exposure. Two formulations were developed to solubilize sertraline at high concentration in the presence of a mucoadhesive.

One of those formulations, containing HPMC as an excipient, successfully maintained stability for up to three months at room temperature and showed promising materials properties.

The formulation was evaluated *in vivo* and was found to be well-tolerated after a single administration in mice. It successfully matched the brain exposure levels of an oral formulation, even though the intranasal administration contained half the dose, thereby demonstrating the enhanced efficiency of intranasal delivery. At the same time, the intranasally treated group showed lower Cmax in the plasma (p = 0.27), suggesting lower systemic exposure.

In contrast, work by another group used a thermo-gelling formulation of sertraline to evaluate the behavioral effects of intranasal administration of sertraline in mice.^45^ The results of this study were surprising and did not align with the data collected here or previously reported pharmacokinetic data for sertraline in mice.^46^ Notably, they found surprisingly low initial plasma concentrations following intravenous administration of sertraline, and the authors concluded that intranasal sertraline did not improve selectivity for the brain, although they suggest that it did enhance behavioral efficacy.^45^ Thus, it is possible that the specific formulation of a drug, in addition to the properties of the drug itself, may impact the resultant biodistribution.

Future work should further investigate the effect of novel intranasal formulations on the selectivity of sertraline for the brain compared to the plasma. Antenatal depression represents a critical unmet need, and a formulation minimizing systemic exposure would be a highly desirable treatment option. If successful, this approach should be applied to other drug classes to mitigate systemic exposure to a wider variety of psychotropic medications for clinical indications such as schizophrenia, obsessive-compulsive disorder, bipolar disorder, and panic disorders. Additionally, it should be explored for the treatment of patients for whom such medications are ineffective due to rapid first-pass metabolism, as intranasal administration has the capacity to overcome this limitation.

## Supporting information

SupportingInformation

## ASSOCIATED CONTENT

The following file is available free of charge:

Supporting Information: experimental methods and supporting data (PDF)

## AUTHOR INFORMATION

### Author contributions

The study was conceptualized by S.C.W. and E.A.A with contributions from V.D. and T.L. Experiments were conducted by S.C.W., T.L., V.D., A.A., D.R.M, C.K.J., and N.E. The original draft was written by S.C.W.

## ACKNOWLEDGEMENTS

The authors thank the SPARK advisors and participants as well as Dr. Emily Meany for their advice and feedback throughout this project.

