## SupportingInformation for "Intranasal sertraline for the investigation of nose-to-brain delivery to mitigate systemic exposure"

**Supplemental methods:**

*Materials*

Sertraline hydrochloride, isopropanol, ethanol, and 1-methyl-2-pyrrolidone were purchased from Fisher Scientific. β-cyclodextrin, dimethyl sulfoxide, glycerol, and hydroxypropyl methylcellulose (US Pharmacopeia Grade) were purchased from Sigma-Aldrich. Pectin (Splendid specialty type 100) was generously provided by CPKelco. The sertraline ELISA kit was purchased from Neogen Corporation.

All absorbance measurements were collected using a Thermo Scientific Nanodrop One. Viscosity measurements were collected using a Rheosense m-VROC viscometer.

*Preparation of formulations*

In order to encapsulate sertraline in β-cyclodextrin, sertraline hydrochloride was dissolved in water with β-cyclodextrin in water at a 1:1 molar ratio. 1 M NaOH was added dropwise to raise the pH and deprotonate sertraline hydrochloride to form the free base of sertraline, which resulted in the formation of a fine white precipitate. The pH was monitored by pH strips. The mixture was heated to 50 °C and stirred until it appeared clear, suggesting the encapsulation of sertraline in β-cyclodextrin. It was then flash-frozen in liquid nitrogen and dried using a Labcono lyophilizer.

To prepare the pectin formulation, the sertraline/β-cyclodextrin powder was dissolved at a concentration of 142 mg/mL (corresponding to ~20 mg/mL of the free base sertraline) in a solvent mixture containing water (53 vol%), glycerol (30 vol%), NMP (12 vol%), and DMSO (5 vol%). The mixture was vortexed until dissolved, and the solution was clear and colorless. This mixture was combined with an equal volume of an aqueous solution containing 2 wt% pectin to yield the final pectin formulation, containing 1 wt% pectin and ~10 mg/mL free base sertraline.

The HPMC intranasal formulation was prepared by dissolving sertraline hydrochloride at a concentration of 26.67 mg/mL in a solvent mixture containing water (53 vol%), glycerol (30 vol%), NMP (12 vol%), and DMSO (5 vol%). The solution was mixed by vortex until it appeared clear and colorless. The mixture was then combined at a 3:1 volume ratio with an aqueous solution of 2 wt% HPMC to yield the final HPMC formulation, containing 0.5 wt% HPMC and 20 mg/mL sertraline hydrochloride.

For the in vitro studies, an additional formulation of HPMC was prepared for oral dosing, which was prepared by the same protocol as the intranasal formulation, but it contained 2.5 wt% sucrose to increase the tolerability for the mice.

*Solubility and stability studies*

The concentration of sertraline in each formulation was measured by monitoring the absorbance at 280 nm, which was then compared to a sertraline standard curve. Formulations were stored at room temperature for the aging assay.

*Viscosity measurements*

Viscosity was measured at 37 °C to mimic biological temperature, which would be relevant after administration. A shear rate of 10000 $s^{-1}$ was implemented, and data were collected after an equilibration period of 2 s.

In vivo *evaluation*

Female C57BL/6 mice ($n$=85), age 8 weeks, were sourced from Charles River Laboratories and cared for according to Institutional Animal Care and Use guidelines. All animal studies were performed in accordance with the National Institutes of Health guidelines and the approval of the Stanford Administrative Panel on Laboratory Animal Care.

The HPMC formulation was selected for *in vivo* evaluation and was freshly prepared within one week of dosing. The sertraline solutions were prepared in sterile Eppendorf tubes and were heated to 80 °C for 1 h to promote sterility before use. Mice were dosed orally with 20 µL or intranasally with 10 µL, for a total dose of 0.4 mg or 0.2 mg. These doses were selected to roughly correlate to a human oral dose of 100 mg daily. Mice were sacrificed by carbon dioxide asphyxiation followed by cardiac puncture at 15 min, 30 min, 2 h, and 4 h after dosing ($n$=9-11 mice per group per timepoint). The brain tissue was harvested and flash-frozen in liquid nitrogen. The lungs, trachea, and nasal passages were collected from three animals of each treatment, sacrificed after 4 h. The samples were collected and preserved for histology by the Stanford Veterinary Services Center Necropsy Lab. The histology slides were prepared with H&E staining by Histo-Tec, and the results were interpreted by a histologist at Stanford Veterinary Services Center Necropsy Lab.

*Sertraline quantification in brain and plasma*

Sertraline concentrations in the brain and plasma were quantified by direct competitive ELISA. Plasma samples were diluted 1:20 in the provided EIA buffer and used directly. Whole brain tissue was homogenized using a glass tissue grinder, using 1 mL of water and 1 mL of the provided EIA buffer per 1 g of brain tissue. The homogenate was centrifuged at 10000 xg for 10 minutes at 4 °C. The supernatant was further diluted 1:5 in the provided EIA buffer before use. Standard calibration curves were prepared by spiking naïve plasma and brain homogenate with a known concentration of sertraline hydrochloride and following the same sample preparation protocol. A one sample t test was performed using GraphPad Prism software to perform the statistical analysis.

**Table S1: Sertraline concentrations in brain and plasma, 15 min** after administration in mice

| Intranasal | | Oral | |
|---|---|---|---|
| **$[sertraline]_{brain}$ (ng/g)** | **$[sertraline]_{plasma}$ (ng/mL)** | **$[sertraline]_{brain}$ (ng/g)** | **$[sertraline]_{plasma}$ (ng/mL)** |
| 562 | 702 | 1085 | 596 |
| 0 | 543 | 0 | 225 |
| 2419 | 1227 | 10793 | 2153 |
| 7226 | 1158 | 5337 | 1469 |
| 4733 | 2796 | 10269 | 1577 |
| 20774 | 4352 | 930 | 534 |
| 12992 | 2408 | 4695 | 1196 |
| 4771 | 2150 | 11370 | 2851 |
| 3249 | 803 | 6928 | 1208 |
| 15164 | 1071 | 1079 | 301 |

**Table S2: Sertraline concentrations in brain and plasma, 30 min** after administration in mice

| Intranasal | | Oral | |
|---|---|---|---|
| **[sertraline]$_{brain}$ (ng/g)** | **[sertraline]$_{plasma}$ (ng/mL)** | **[sertraline]$_{brain}$ (ng/g)** | **[sertraline]$_{plasma}$ (ng/mL)** |
| 2987 | 1666 | 7952 | 2320 |
| 3892 | 2130 | 5435 | 1493 |
| 8872 | 2581 | 6744 | 711 |
| 10177 | 2333 | 11879 | 1121 |
| 5190 | 623 | n.d. | 1402 |
| 4864 | 1170 | 12136 | 4896 |
| 2282 | 1141 | 4569 | 1449 |
| 11485 | 2371 | 7664 | 2692 |
| 5589 | 1113 | 2199 | 1661 |
| 7422 | 2299 | 36068 | 5513 |
| 16702 | 1365 | | |

**Table S3: Sertraline concentrations in brain and plasma, 2 h** after administration in mice

| Intranasal | | Oral | |
|---|---|---|---|
| **$[\text{sertraline}]_{\text{brain}}$ (ng/g)** | **$[\text{sertraline}]_{\text{plasma}}$ (ng/mL)** | **$[\text{sertraline}]_{\text{brain}}$ (ng/g)** | **$[\text{sertraline}]_{\text{plasma}}$ (ng/mL)** |
| 8678 | 5258 | 12588 | 521 |
| 14485 | 1730 | 7904 | 4747 |
| 7168 | 1187 | 12362 | 2380 |
| 9627 | 631 | 11272 | 30806 |
| 4192 | 779 | 5890 | 1241 |
| 7484 | 2008 | 6455 | 1109 |
| 9173 | 1427 | 7880 | 1327 |
| 2284 | 1521 | 11183 | 1280 |
| 7355 | 1957 | 4230 | 1291 |
| 9591 | 430 | 4168 | 930 |

**Table S4: Sertraline concentrations in brain and plasma, 4 h** after administration in mice

| Intranasal | | Oral | |
|---|---|---|---|
| **$[sertraline]_{brain}$ (ng/g)** | **$[sertraline]_{plasma}$ (ng/mL)** | **$[sertraline]_{brain}$ (ng/g)** | **$[sertraline]_{plasma}$ (ng/mL)** |
| 15379 | 1514 | 10076 | 783 |
| 11267 | 1009 | 5762 | 775 |
| 4104 | 642 | 6670 | 417 |
| 6283 | 1190 | 13571 | 1431 |
| 8143 | 1903 | 8295 | 1942 |
| 6804 | n.d. | 7063 | 1104 |
| 8363 | 1399 | 6786 | 434 |
| 5493 | 927 | 5117 | 735 |
| 3343 | 289 | 3897 | 579 |
| 7875 | 174 | 13272 | 1019 |

**Table S5: Results of histological analysis**, as reported by Stanford Necropsy Lab

| Treatment | Body Weight (g) | Brain Weight (g) | Gross Notes | Trachea | Thyroid | Thymus | Hilar Lymph Node | Lung | Nasal Cavity |
|---|---|---|---|---|---|---|---|---|---|
| IN (0.2 mg) | 20.5 | 0.4437 | WNL | WNL; rare PMNs at bifurcation | WNL | WNL | WNL | WNL | Rare PMN's in lamina propria; small ulcerations; ventral rostral portion |
| PO (0.4 mg) | 15.8 | 0.4171 | Malocclusion lower incisors | WNL | WNL | --- | --- | WNL | Rare PMN's in lamina propria; small ulcerations; ventral rostral portion |
| PO (0.4 mg) | 19.8 | 0.4516 | WNL | WNL | WNL | --- | --- | WNL | Rare PMN's in lamina propria; ventral rostral portion |
| PO (0.4 mg) | 20.3 | 0.4614 | WNL | *Mainstem bronchi; rare PMNs at bifurcation | --- | WNL | WNL | WNL | Rare PMN's in lamina propria; ventral rostral portion and VNO |
| IN (0.2 mg) | 20.1 | 0.4413 | WNL | WNL | WNL | WNL | WNL | WNL | Rare PMN's in lamina propria; small ulcerations; ventral rostral portion |
| PO (0.4 mg) | 17.9 | 0.4213 | WNL | WNL | WNL | --- | --- | WNL | Rare PMN's in lamina propria; small ulcerations; ventral rostral portion |
